# Using a stable protein scaffold to display peptides that bind to alpha-synuclein fibrils

**DOI:** 10.1101/2025.02.04.636406

**Authors:** Samuel Bismut, Matthias M. Schneider, Masashi Miyasaki, Yuqing Feng, Ellis J. Wilde, M. Dylan Gunawardena, Tuomas P.J. Knowles, Gabrielle S. Kaminski Schierle, Laura S. Itzhaki, Janet R. Kumita

**Affiliations:** Department of Pharmacology, University of Cambridge, Tennis Court Road, Cambridge, UK, CB2 1PD; Yusuf Hamied Department of Chemistry, University of Cambridge, Lensfield Road, Cambridge, UK, CB2 1EW; Department of Chemical Engineering and Biotechnology, University of Cambridge, Cambridge, UK, CB3 0AS

**Author notes:** Correspondence: Janet Kumita, Department of Pharmacology, University of Cambridge, Cambridge UK, CB2 1PD. These authors contributed equally (Samuel Bismut, Matthias M. Schneider).

**Keywords:** Amyloid fibrils, α-synuclein, protein scaffolds, fibril-binders

## Abstract

Amyloid fibrils are ordered aggregates that are a pathological hallmark of many neurodegenerative disorders including Alzheimer’s Disease and Parkinson’s Disease. The process of amyloid formation involves a complex cascade by which soluble monomeric protein converts to insoluble, ordered aggregates (amyloid fibrils). Although inhibiting the aggregation pathway is a key target for therapeutic development, the heterogeneous collection of aggregation-prone species formed in this process, including oligomers, protofibrils and fibrils, represent other targets for modifying disease pathology. Developing molecules that can bind to amyloid fibrils and potentially disrupt the harmful interactions between the fibrils and the cellular components would be advantageous. Designing peptide modulators for α-synuclein aggregation is of great interest, however effective inhibitory peptides are often hydrophobic and hence difficult to handle; therefore, developing strategies to display these peptides in a soluble scaffold would be very beneficial. Here we demonstrate that the ultra-stable consensus-designed tetratricopeptide repeat (CTPR) protein scaffold can be grafted with “KLVFF” derived peptides previously identified to inhibit protein aggregation and interact with amyloid fibrils, to produce proteins that bind along the surface of α-synuclein fibrils with micromolar affinity. Given the ability to insert hydrophobic peptides to produce soluble, CTPR-based binders, this method may prove beneficial in screening for peptide modulators of protein aggregation.

**Short statement:** Consensus-designed tetratricopeptide repeat (CTPR) proteins can be endowed with additional functionality including specific client recruitment. Here we created stable, soluble CTPR-based binders that recognise α-synuclein fibrils with micromolar affinity. The CTPR scaffold provides a facile way to test potential fibril-binding or aggregation-modulating peptides and to probe their mechanisms of action.

## Introduction

Protein misfolding and aggregation into amyloid fibrils has been identified as a pathological hallmark for several neurodegenerative disorders, including Alzheimer’s Disease (AD) and Parkinson’s Disease (PD) (Chiti and Dobson 2017). In PD, the 14 kDa protein α-synuclein aggregates to form cross-β amyloid fibrils via a pathway that populates less organized oligomeric intermediates (Li et al. 2001; Serpell et al. 2000). Although it is believed that these oligomeric species are the key toxic species in the context of disease (Cremades et al. 2012; Fusco et al. 2017; Glabe 2006), the mature fibrils have a key role in propagating the aggregation process, for example, through secondary nucleation processes where protofibrils can accelerate amyloid formation through their fibril ends or fibril-surfaces (Buell et al. 2014; Cohen et al. 2013). Along with significant efforts to identify inhibitors of the aggregation process of α-synuclein (Agerschou et al. 2019; Cheruvara et al. 2015; Chia et al. 2023; Meade et al. 2020; Santos et al. 2021), there has been great interest in identifying molecules that can bind specifically to the α-synuclein fibrils, in order to modulate the aggregation process (Monsellier et al. 2020; Sangwan et al. 2020; Scheidt et al. 2021; Wallace et al. 2024). Furthermore, fibril-binding strategies are important for targeted protein degradation approaches to direct aggregates to the cell’s natural proteostasis machinery for clearance (Lee et al. 2023; Tomoshige and Ishikawa 2021). Recent advances in *de novo* design have enabled the development of peptides and mini-proteins that bind to amyloid fibrils or with regions of monomeric amyloid-related proteins, creating high-affinity scaffolds that can modulate aggregation pathways (Agerschou et al. 2019; Sahtoe et al. 2024; Sangwan et al. 2020; Wallace et al. 2024). Interestingly, peptides derived from the KLVFF peptide (amino acids 16-20; Aβ_1-42_) that is known to modulate Aβ fibril formation (Tjernberg et al. 1996), have also been reported to bind to Aβ fibrils and α-synuclein fibrils (Aoraha et al. 2015; Wood et al. 2020). There are several studies that use the KLVFFAE peptide to bind to Aβ fibrils, but the peptide itself is aggregation-prone and consequently often requires methods to increase its solubility and efficacy, including terminal-capping strategies (Aoraha et al. 2015), incorporating D-amino acids (Horsley et al. 2020), and coupling to gold and gadolinium nanoparticles (Gao et al. 2015; Plissonneau et al. 2016).

To test potential fibril-binding peptides and probe the mechanism by which they interact with amyloid fibrils, having an ultrastable scaffold for their display would be advantageous. The consensus-designed tetratricopeptide repeat protein (CTPR) scaffold comprises tandem arrays of a 34-residue α-helix-turn-α-helix motif, and their simple, modular architecture and high stability makes them useful tools for protein engineering (Main et al. 2003; Perez-Riba and Itzhaki 2019). A rational design approach can be used to endow CTPR scaffolds with additional functionality including specific client recruitment via grafting of single and multiple copies of short linear binding motifs (SLiMs) between adjacent repeats in CTPR scaffolds (ranging from 2-6 repeat units), with diverse, precise, and predictable geometries for multivalent and multi-functional display (Diamante et al. 2021; Madden et al. 2019; Ng 2024; Perez-Riba and Itzhaki 2019). Here we demonstrate that the KLVFFAE peptide and a variation, KLVFWAK (Wood et al. 2020), can be grafted onto our 3-repeat CTPR scaffold to produce soluble chimeras (FibSyn-CTPR and FibSyn2-CTPR) that can bind with micromolar affinity to α-synuclein fibrils, along with a 4-repeat version containing two KLVFFAE motifs (between repeat 1-2 and between repeat 3-4; 2XFibSyn-CTPR4) that displays similar affinity for α-synuclein fibrils with an increase in the number of associated molecules to the fibrils. Incubation of fluorophore-labelled fibril-binding CTPRs with unlabelled α-synuclein fibrils resulted in full visualisation of the fibrillar structures by fluorescence microscopy, revealing that the interaction is with the entire surface of the fibrils and not isolated to specific regions. Finally, constraining these peptides in the inter-repeat loop of the CTPR-scaffold imparts a preference of the FibSyn-CTPR variants for α-synuclein fibrils as compared to pre-formed Aβ_1-42_ fibrils.

## Results

Single peptide motifs (KLVFFAE or KLVFWAK) were grafted between repeat 1-2 of a CTPR3 scaffold designed with a solvating helix (Main et al. 2003) to create FibSyn-CTPR3 and FibSyn2-CTPR3, respectively. A CTPR4 scaffold containing the KLVFFAE peptide between repeat 1-2 and between repeat 3-4 (2XFibSyn-CTPR4) was also designed (Figure 1). The CTPR4 design does not include a solvating helix, as Diamante and co-workers showed that the CTPR4 scaffold can accommodate 2-grafted binding loops without compromising the solubility or protein yield (Diamante et al. 2021). In all cases, DPNN linker regions flanked the fibril-binding peptides to increase the flexibility of the loops to maximise binding (Madden et al. 2019).

**Figure 1:**
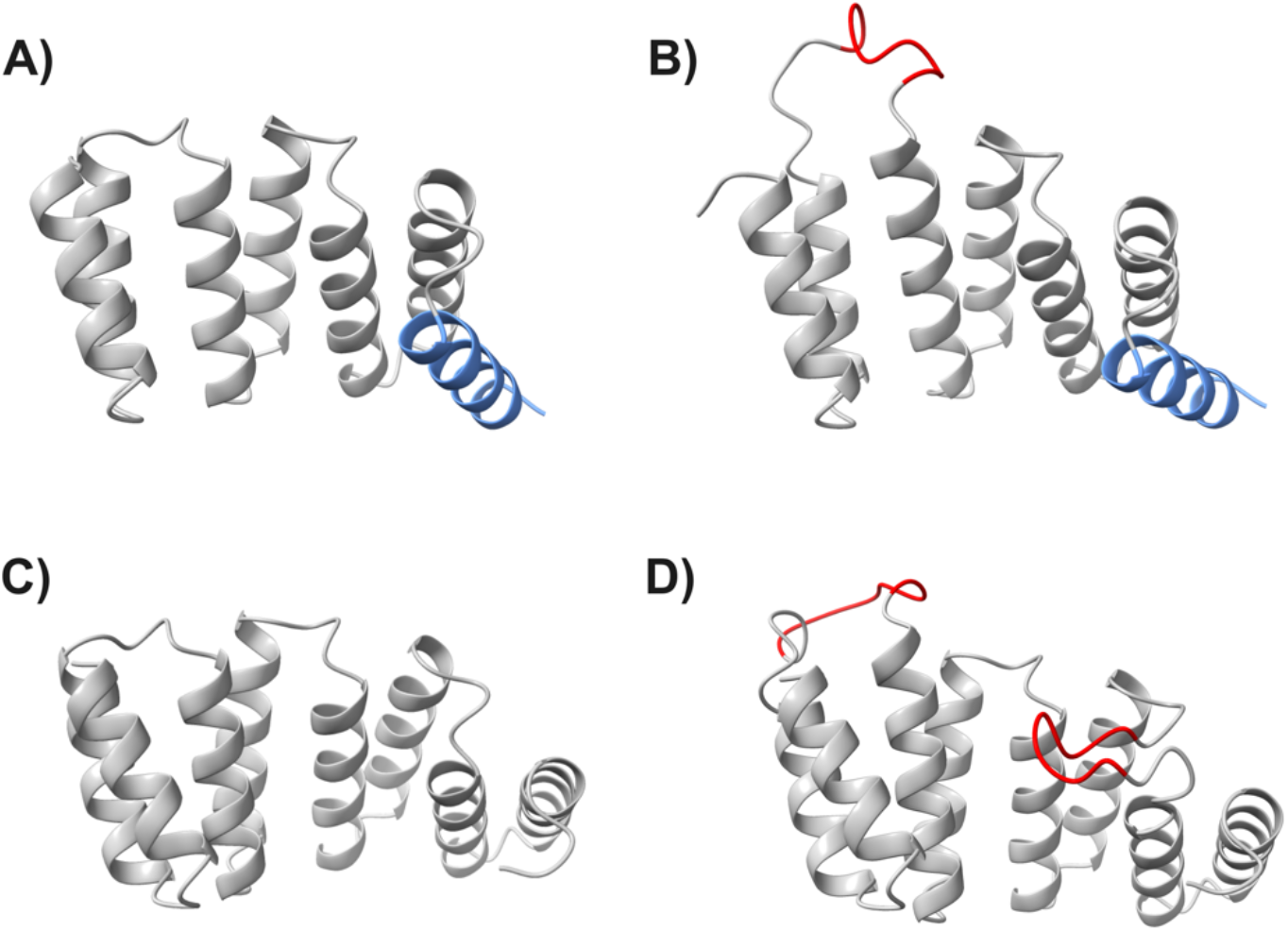
Schematic diagrams of CTPR scaffolds with incorporated fibril-binding motifs. Alphafold2 models of A) the CTPR3 scaffold protein containing three-repeat units and a solvating helix (blue), B) FibSyn-CTPR3 containing three-repeat units, a solvating helix and a fibril-binding motif between repeat 1-2 (red), C) CTPR4 containing four-repeat units and D) 2XFibSyn-CTPR4 consisting of four-repeat units with a fibril-binding motif between repeat 1-2 and between repeat 3-4 (red). The peptide sequence for FibSyn is KLVFFAE and for FibSyn2 it is KLVFWAK. Full protein sequences are available in Table S1.

### Fibril-binding CTPRs retain scaffold structure and high stability

All CTPR variants (and their respective negative controls wild-type (WT) CTPR3 and CTPR4, containing no additional inter-repeat loops) were expressed and purified, resulting in protein concentrations in the range of 120-320 µM. The insertion of the fibril-binding loops does not alter the native fold of the CTPR variants compared with their WT controls, as measured by circular dichroism (CD) spectroscopy (Figure 2, Fig. S1), and as expected, they decrease the native-state stability as determined by GdnHCl denaturation assays, but only by a modest amount (Table 1, Fig. S1, S2). In agreement with the chemical denaturation, thermal unfolding, assayed by measuring the CD spectrum at 90°C, shows that the helical content of the CTPR variants decreases more than those of the WT controls (Fig. S2) but the helicity is recovered when returning to 20°C (Figure 2).

**Table 1.** Parameters obtained from two-state fit of the GdnHCl-induced chemical denaturation experiments for CTPR variants.

| Protein | $D_{50\%}$ (M) | $m$-value ($\text{kcal mol}^{-1} \text{M}^{-1}$) |
| --- | --- | --- |
| CTPR3 | $3.6 \pm 0.1$ | $2.6 \pm 0.2$ |
| FibSyn-CTPR3 | $3.4 \pm 0.1$ | $2.1 \pm 0.5$ |
| FibSyn2-CTPR3 | $2.8 \pm 0.2$ | $1.3 \pm 0.7$ |
| CTPR4 | $4.8 \pm 0.1$ | $3.8 \pm 0.6$ |
| 2XFibSyn-CTPR4 | $3.8 \pm 0.1$ | $1.4 \pm 0.5$ |

**Figure 2:**
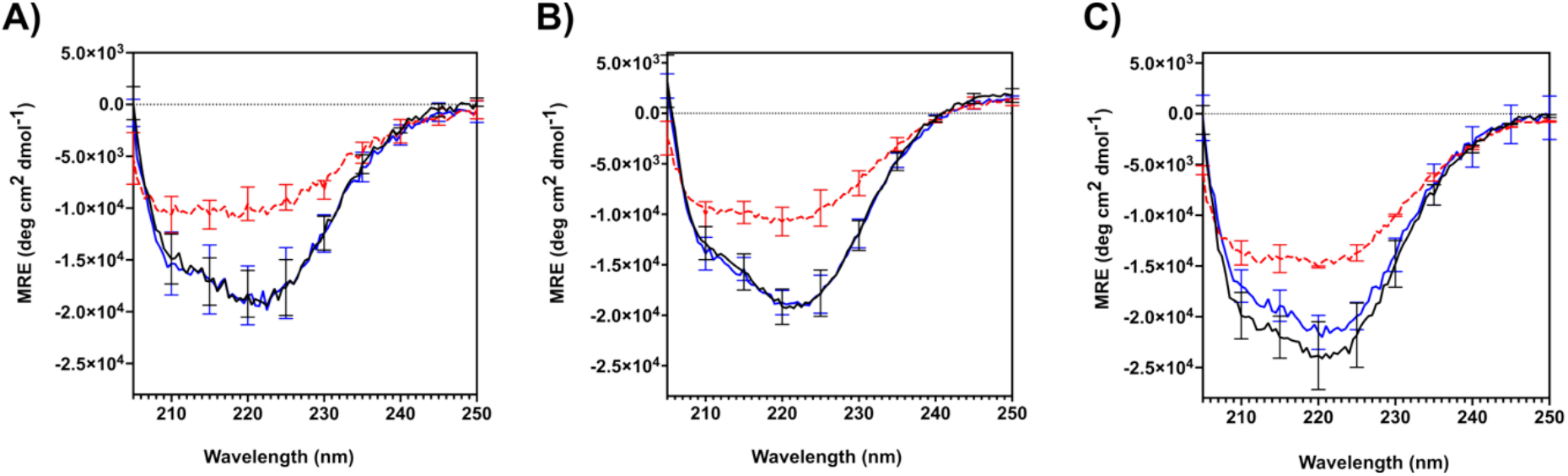
CD analysis of CTPR variant secondary structure. Far-UV CD spectra of A) FibSyn-CTPR3, B) FibSyn2-CTPR3 and C) 2XFibSyn-CTPR4 at 20°C (black), 90°C (red) and returning to 20°C (blue). Protein concentration was 5 µM in 50 mM sodium phosphate pH 7.5, 150 mM NaCl. Error bars are standard deviation calculated from three independent experiments.

### Fibril-binding CTPR variants interact with α-synuclein fibrils

To test whether the CTPR variants with inserted peptide motifs could recognize monomeric or fibrillar α-synuclein, an initial dot blot assay was performed. After applying monomeric and fibrillar α-synuclein to nitrocellulose membranes, fluorescently labelled CTPR variants were incubated to look for interactions. FibSyn-CTPR3, FibSyn2-CTPR3 and 2XFibSyn-CTPR4 showed positive interactions with α-synuclein fibrils but not monomeric α-synuclein (Fig. S3) and CTPR3-WT showed little interaction. To ensure that these specific interactions were not an artifact of immobilising the fibrils onto the nitrocellulose membrane, a fibril-pulldown assay was carried out. CTPR variants containing a single cysteine were labelled with Alexa Fluor^TM^ 647 maleimide (Figure 3A) and incubated with α-synuclein fibrils (Figure 3B). After extensive washing of the fibril/CTPR complexes to remove non-specific fluorescently labelled CTPRs, the fluorescence emission spectra were measured (Figure 3C, D) and corrected for labelling efficiency. Using these experiments, we could observe clear binding between the FibSyn-CTPR3, FibSyn2-CTPR3 and 2XFibSyn-CTPR4 variants and α-synuclein fibrils. We observed lower fluorescence intensity for CTPR3-WT (Figure 3C) indicative of weaker binding, which we also noted in the dot-blot assay, whereas CTPR4-WT showed little fluorescence emission.

**Figure 3:**
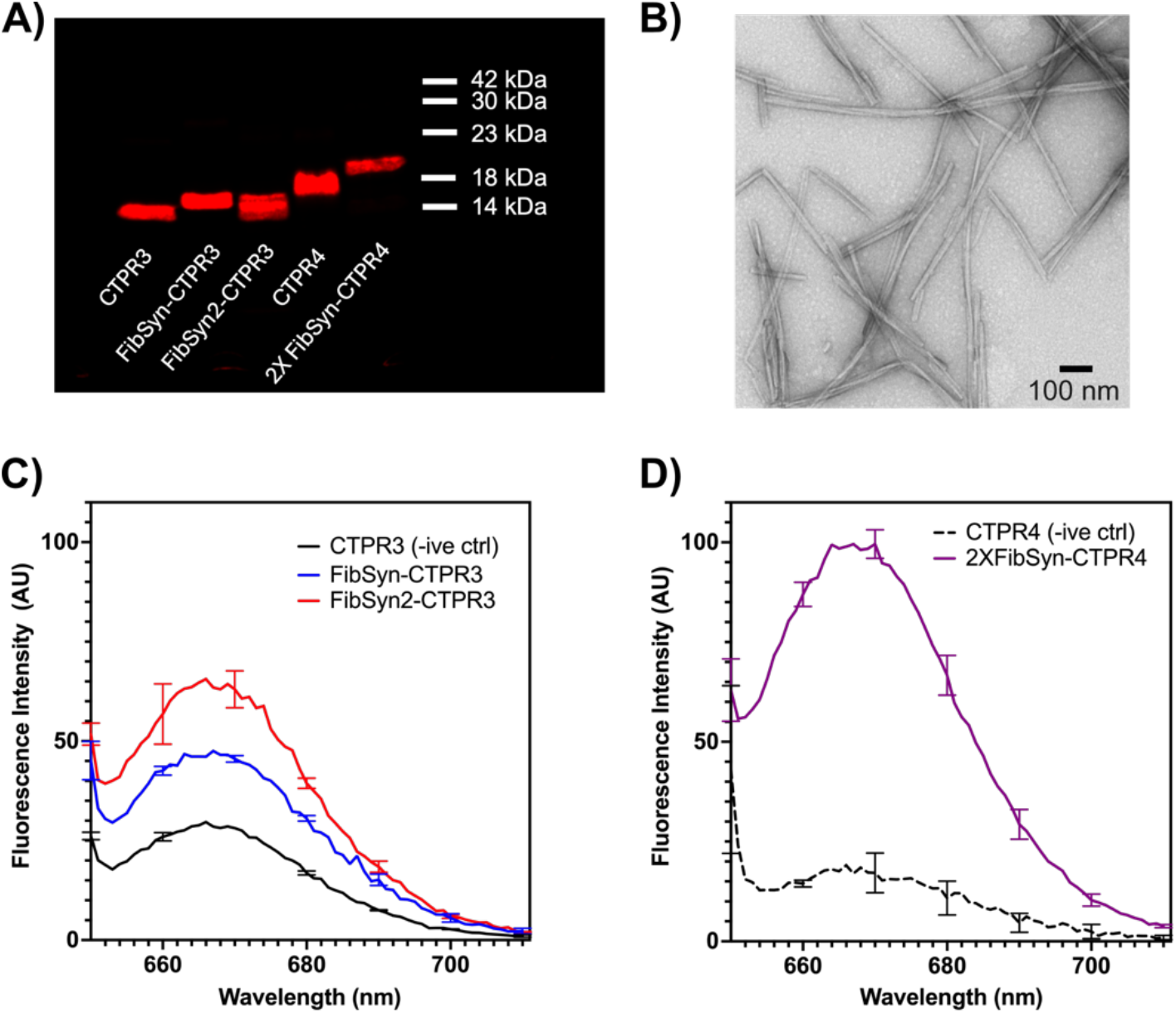
Fibril pulldown assay to assess interactions between CTPR variants and α-synuclein fibrils. A) SDS-PAGE analysis of Alexa Fluor^TM^ 647 labelled CTPR variants. B) TEM image of α-synuclein fibrils prior to incubation with fluorescently labelled CTPRs. C) Fluorescence emission spectrum for CTPR3 (negative control), FibSyn-CTPR3 and FibSyn2-CTPR3 bound α-synuclein fibrils. D) Fluorescence emission spectrum for CTPR4 (negative control) and 2XFibSyn-CTPR4 bound α-synuclein fibrils. Error bars represent the standard deviation of two independent experiments.

### Fibril-binding CTPR variants bind with micromolar affinity and along the entire fibril surface

To measure the binding affinities and stoichiometries of the interactions between the fibril-binding CTPRs and α-synuclein fibrils, we applied a microfluidic diffusional sizing (MDS) approach. MDS allows us to determine the hydrodynamic radii (R_H_) of individual components in complex mixtures of species and to quantify their relative concentrations (Arosio et al. 2016; Scheidt et al. 2021; Scheidt et al. 2019). Using fluorescently labelled CTPR variants, we can monitor the change in R_H_ as they bind to large, pre-formed fibrils (PFFs) of α-synuclein to quantify these interactions. MDS also provides insights into the stoichiometry of the CTPR variants in relation to the α-synuclein units.

An initial comparison of the hydrodynamic radius of Alexa Fluor^TM^ 647-labelled FibSyn-CTPR3 with increasing concentrations of unlabelled α-synuclein PFFs shows a significant increase in R_H_, whereas the same titration with Alexa Fluor^TM^ 647-labelled CTPR3-WT shows no binding (Figure 4A). Next, we performed titrations for each fibril-binding CTPR variant and by analyzing the fraction of bound CTPR-variant at varying PFF concentrations, we determined the dissociation constants and stoichiometries (Table 2). All three fibril-binding variants had dissociation constants in the low micromolar range. When comparing the stoichiometry, i.e. the number of CTPR binders per α-synuclein unit, the FibSyn-CTPR3 and FibSyn2-CTPR3 were very similar, 1 binder per 34 and 1 binder per 45, respectively, whereas for 2XFibSyn-CTPR4 the stoichiometry was approximately doubled with 1 binder per 14 α-synuclein units. This result suggests that, although the presence of two KLVFFAE peptide motifs does not increase the affinity of the CTPR variant for the fibrils, both binding motifs interact with the fibrils. Finally, to determine the localization of the fibril-binding CTPR variants on α-synuclein fibrils, we imaged the Alexa Fluor^TM^ 647-labelled CTPR variants bound to α-synuclein fibrils using total internal reflection fluorescence (TIRF) microscopy (Figure 4E). In all cases, the CTPR variants are distributed throughout the fibril structures.

**Table 2:**
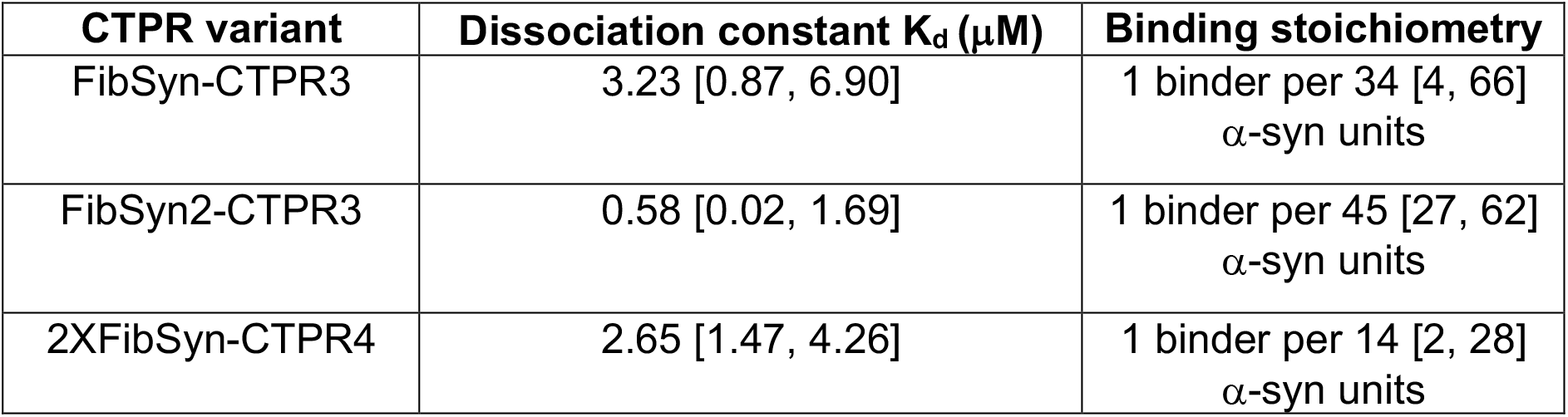
Binding of CTPR variants with α-synuclein PFFs. Values calculated from MDS experiments with the upper and lower boundaries of the 95% confident interval shown in square brackets.

| **CTPR variant** | **Dissociation constant $K_d$ (μM)** | **Binding stoichiometry** |
| --- | --- | --- |
| FibSyn-CTPR3 | 3.23 [0.87, 6.90] | 1 binder per 34 [4, 66]<br>α-syn units |
| FibSyn2-CTPR3 | 0.58 [0.02, 1.69] | 1 binder per 45 [27, 62]<br>α-syn units |
| 2XFibSyn-CTPR4 | 2.65 [1.47, 4.26] | 1 binder per 14 [2, 28]<br>α-syn units |

**Figure 4:**
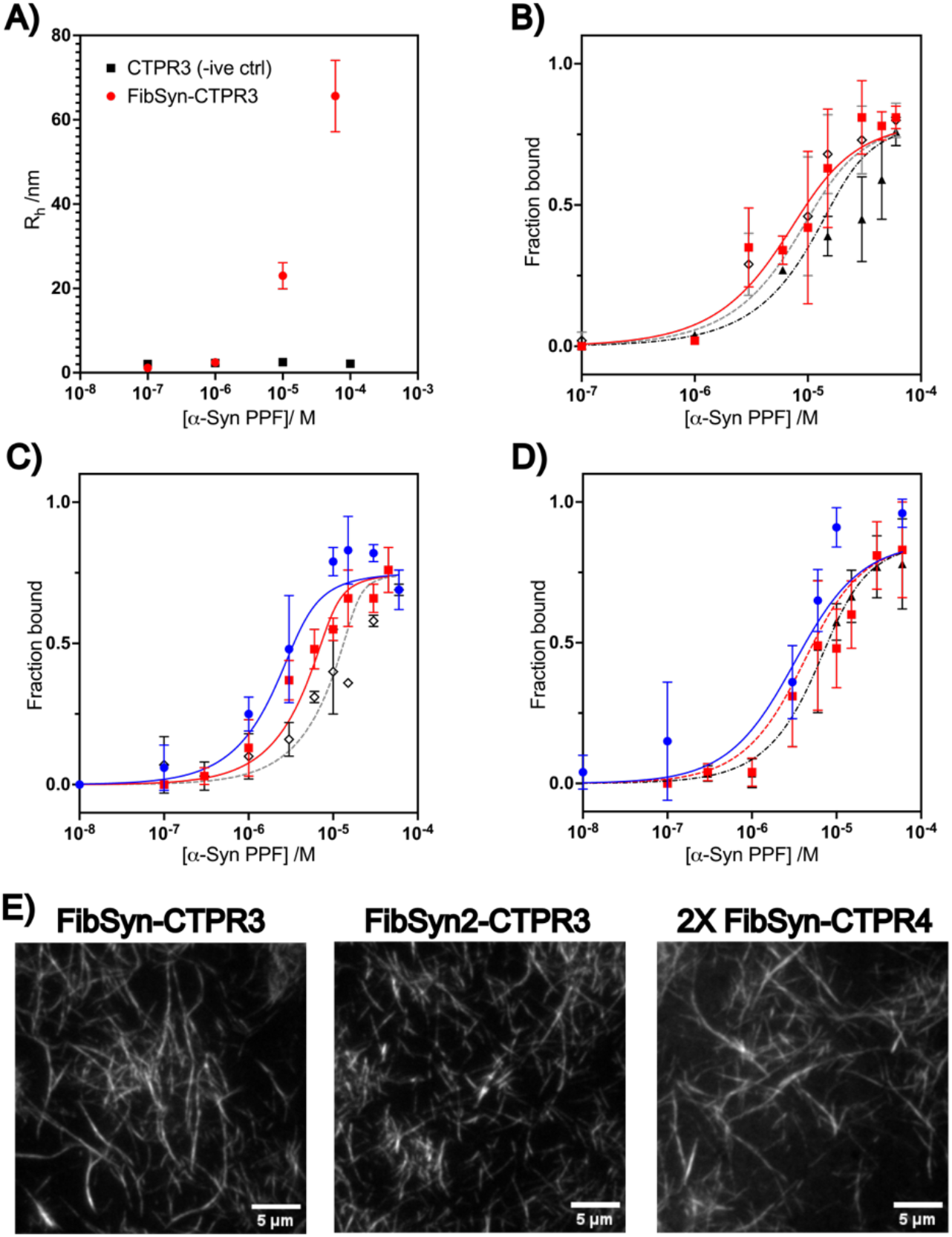
Microfluidic diffusional sizing (MDS) assay to determine binding affinity between fluorescently labelled CTPR variants and α-synuclein preformed fibrils (PFFs). A) Initial hydrodynamic radius measurements to compare CTPR3 (negative control) and FibSyn-CTPR3 interactions with increasing α-synuclein PFF concentrations. The hydrodynamic radius for FibSyn-CTPR3 significantly increases in the presence of α-synuclein PFFs whereas no change is observed for CTPR3 (negative control). Fraction of FibSyn bound to α-synuclein PFFs at varying concentrations of PFFs for B) FibSyn-CTPR3, C) FibSyn2-CTPR3 and D) 2XFibSyn-CTPR4. Each MDS assay was performed with different concentrations of CTPR variants including 75 nM (blue circles), 200 nM (red squares), 400 nM (black open diamonds) and 500 nM (black triangles). Dissociation constants are shown in Table 2. E) TIRF microscopy imaging of Alexa Fluor^TM^ 647-labelled CTPR variants bound to α-synuclein fibrils (prepared in a similar way to Figure 3 pulldown assay samples). Scale bars represent 5 µm.

As the KLVFFAE peptide motif is well studied for its interactions with amyloid-β (Aβ) aggregates (Aoraha et al. 2015; Liu et al. 2023; Plissonneau et al. 2016; Wood et al. 2020), we tested if our fibril-binding CTPR variants showed specific interactions with pre-formed Aβ_1-42_ fibrils, using first a fibril pulldown assay for FibSyn2-CTPR3 versus CTPR3-WT and followed by MDS experiments for all three potential binders. In both experiments, we observe no interactions between the fibril-binding CTPR variants and Aβ_1-42_ fibrils (Fig. S4). It appears that constraining this peptide-motif within the inter-repeat loops of the CTPR scaffold imparts a conformational restraint that allows these variants to have a specificity for α-synuclein fibrils over Aβ_1-42_ fibrils.

## Discussion

Here we show the utility of the CTPR scaffold to not only display hydrophobic peptides in the inter-repeat loops to produce highly soluble protein, but to also produce CTPR-binders that have micromolar affinity for α-synuclein fibrils. In addition, these CTPR-binders can be used to label the surfaces of *in vitro* α-synuclein to enable visualization of the structures using fluorescence microscopy. It is interesting to note that constraining these peptides in the inter-repeat loops of the CTPRs introduces a selectivity for α-synuclein fibrils over Aβ_1-42_ fibrils that has not been reported for the peptide motifs alone (Aoraha et al. 2015; Wood et al. 2020). Despite the similarities in the cross-β structure of amyloid fibrils, the physicochemical attributes can differ significantly, and this may impact the fibril-binding properties of our CTPR-binders. For example, spectrally-resolved point accumulation for imaging in nanoscale topography (sPAINT) microscopy techniques have demonstrated that the surfaces of *in vitro* α-synuclein fibrils are more hydrophobic than Aβ fibrils (Bongiovanni et al. 2016). However, further studies are needed to elucidate the binding epitopes on the α-synuclein fibrils that are recognized by these CTPR-binders.

Although the interaction of the FibSyn-CTPRs with the surfaces of the synuclein fibrils may have implications for modulating aggregation kinetics (Buell et al. 2014), attempts to establish the effect on surface-catalyzed nucleation assays have been inconclusive, in large part due to the kinetic reaction conditions being carried out at a pH that is not ideal for the FibSyn-CTPRs, following close to the isoelectric points. Within the CTPR scaffold, only 8 residues are essential to ensure correct folding and, therefore, this system is amenable to protein engineering to improve its physicochemical properties (Main et al. 2003; Uribe et al. 2021); rational design approaches may lead to future variants that can be used to modulate aggregation kinetics.

Given the number of peptides reported in literature that alter α-synuclein aggregation or bind to different α-synuclein conformers (Cheruvara et al. 2015; Meade et al. 2020; Meade et al. 2021; Monsellier et al. 2020; Nahomi et al. 2015; Santos et al. 2021; Wood et al. 2020), our CTPR scaffold may provide a facile method to further study their mechanisms of action for modulating amyloid fibril formation and lead to the development of diagnostic tools and potential therapeutics.

## Material and Methods

### CTPR plasmid preparation and protein purification

CTPR constructs were cloned using gBlock oligos (Integrated DNA technologies) into the pRSET-B vector with FastDigest BamHI and HindIII restriction enzymes (ThermoFisher Scientific (UK) Ltd.) and Anza T4 DNA ligase master mix (Invitrogen, Paisley, UK). Plasmids were transformed into chemically competent C41 *E. coli* cells using standard heat-shock protocols. Colonies were grown at 37°C in 2xYT media (Formedium, Swaffham, UK) containing ampicillin (50 µg/mL). Upon reaching OD_600nm_ of 0.6-0.8, bacteria cultures were induced with isopropyl D-thiogalactopyranoside (IPTG, 0.5 mM, PanReac AppliChem, Darmstadt, Germany) and incubated (18 h, 20°C, 200 rpm). Cell pellets were collected by centrifugation from 125 mL of culture and resuspended in 15 mL lysis buffer (50 mM sodium phosphate (pH 8.0), 150 mM NaCl, 1 mg/mL DNase I, 1 cOmplete EDTA-free protease inhibitor tablet (Roche Diagnostics GmbH) for each 50 mL of lysis buffer). Cells were lysed using an EmulsiFlex C5 homogeniser (Avestin, Mannheim, Germany) (3-4X, 10000-15000 psi) and centrifuged. The supernatants were purified using HisTrap HP columns (1 mL, Cytiva Ltd., Little Chalfont, UK) on an AKTA PURE protein purification system (Cytiva Ltd.) with Buffer A (50 sodium phosphate (pH 8.0), 150 mM NaCl, and a linear gradient (0-80% Buffer A + 0.5 M imidazole, 15 column volumes (CV)). Purification buffers for the Cys-variants contained 0.5 mM TCEP. Protein purity was confirmed using SDS-PAGE and electrospray ionisation mass spectrometry performed on a Xevo G2 mass spectrometer with data analyzed using MassLynx software (Waters UK) (Yusuf Hamied Department of Chemistry, University of Cambridge, UK). The amino acid sequences and masses of the proteins are listed in (Table S1).

### Alexa Fluor^TM^ 647-maleimide labelling of Cys-CTPR variant

Cys-variants of the CTPR proteins (100 µM, 50 mM sodium phosphate buffer, 150 mM NaCl, pH 8, 0.5 mM TCEP) were labelled by reacting with a 2.5X molar excess of Alexa Fluor^TM^ 647 C_2_ maleimide or Alexa Fluor^TM^ 594 C_2_ maleimide (Invitrogen) and incubated (2 hr, room temperature (RT)). 1 mM DTT was added, and excess dye was removed using Pierce dye removal columns as described in the manufacturer’s protocol (Thermofisher). UV-vis spectra for the fluorophore-labelled protein were collected on a Nanodrop 2000 spectrophotometer (Thermofisher) and the protein concentration and labelling efficiency were calculated using Equations 1 and 2 (Alexa Fluor^TM^ 594) and Equations 3 and 4 (Alexa Fluor^TM^ 647):

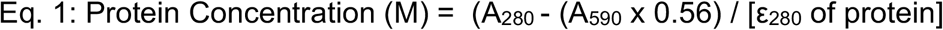

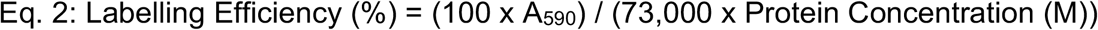

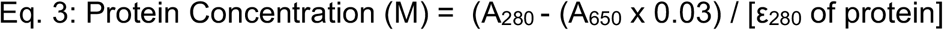

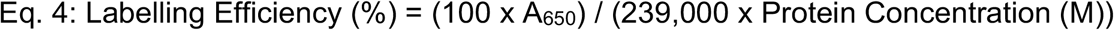

### Circular Dichroism (CD) spectroscopy

CD measurements were conducted with a Chirascan CD spectrometer (Applied Photophysics, Leatherhead, UK) in 1 mm pathlength Precision Cells (110-QS; Hellma Analytics, Müllheim, Germany). Protein samples at 5 µM in sodium phosphate buffer were measured across 205-250 nm wavelengths. Spectra, taken at 1 nm intervals every 0.5 s, were recorded at 20°C, then 90°C, and again at 20°C. Each protein measurement was independently repeated three times, with data baseline corrected for buffer and averaged.

### Guanidine Hydrochloride (GdnHCl) denaturation assay

The different GdnHCl concentrations were prepared by mixing the appropriate volumes of 50 mM sodium phosphate buffer pH 7.5, 150 mM NaCl, 7 M GdnHCl and 50 mM sodium phosphate buffer using a Microlab ML510B (Hamilton, Bonaduz, Switzerland). The protein concentration used was 15 μM and protein and solutions were dispensed into 96-well, half-area, black polystyrene plates (Corning) and covered with 96-well microplate aluminium sealing tape (Corning). Samples were equilibrated at 25 °C for 2 hr. Emission at 360 nm was measured using a CLARIOstar® Microplate Reader (BMG Labtech, Aylesbury, UK). The top optic was used in precise mode for 2 cycles with an excitation wavelength of 295 nm and a dichroic PL325 nm filter at 25°C. Each measurement was performed in triplicate and three independent experiments were carried out for each protein variant unless otherwise specified. The data were fit to a two-state model, with denaturation curves fitted directly in GraphPad Prism 10 as described by Perez-Riba et al. (Perez-Riba and Itzhaki 2017).

### α-Synuclein fibril preparation

A T7-7 plasmid encoding α-synuclein (Hoyer et al. 2002) was transformed and expressed in BL21(DE3) E. coli (NEB (UK) Ltd.), and purification was carried out as detailed in Bell et al. (Bell et al. 2023). α-Synuclein fibrils were generated by incubating 100 µM α-synuclein (PBS with 0.01% azide) for 72-96 hr (37°C, 200 rpm). After incubation, fibril samples were mixed with Thioflavin-T (ThT) (250 µM) and ThT fluorescence was measured to using a CLARIOstar® Microplate Reader (BMG Labtech) with excitation at 440 nm and emission spectra collected between 465-580 nm. The fibrils were centrifuged (10 min, 15,000g) and the supernatant was removed. The fibril pellets were washed with PBS (2X) followed by centrifugation. The washed fibrils were resuspended in PBS and sonicated (low power, 50% pulse, 15 sec) with a Sonopuls probe sonicator (Bandelin, HD 2070) to produce preformed fibrils (PFFs). These PFFs were used to seed fresh α-synuclein fibrils by adding 10% v/v to monomeric protein and incubating for 24-48 hr (37°C, quiescent). All fibril samples were imaged by Transmission electron microscopy (TEM) prior to using. For TEM analysis, fibrils were diluted to 5-10 µM and incubated (3 min) on carbon-coated copper grids (EM Resolutions, Keele, UK), followed by washing with deionised water and staining with UranyLess (Labtech, Rotheram UK) for 2 min (2X) then washed quickly with water. TEM images were taken on a Tecnai G2 80-200 kV transmission electron microscope (Cambridge Advanced Imaging Centre (CAIC), University of Cambridge. Images were analysed using the SIS Megaview II Image Capture system.

### Dot blot assay

Onto nitrocellulose membranes, 5 µL drops of PBS only, α-synuclein monomer (10 µM) and α-synuclein fibrils (10 µM) were applied and dried. This was repeated four times. The membrane was incubated in 2% w/v bovine serum albumin (BSA) in TBS-T (Tris buffered saline with 0.1% v/v Tween 20) (4°C, overnight, roller). The membrane was incubated with a fluorophore-labelled CTPR variant solution (1 µM, 50 mM phosphate pH 8, 150 mM NaCl) (1 hr, RT, roller). The membrane was washed by incubating in TBS-T (3X, 10 min) and then imaged with a LiCor Odyssey (Li-Cor, Lincoln, USA) at 600 nm with a 2 min exposure (Alexa Fluor^TM^ 594) or 700 nm with a 2 min exposure (Alexa Fluor^TM^ 647) and analysed with Empiria Studio^TM^ software (Li-Cor).

### Fibril pulldown assay

Aliquots of α-synuclein fibrils (35 µL of 70 µM solutions (based on monomer concentration)) were incubated with Alexa Fluor^TM^ 647-labelled CTPR variants (35 µL of 20 µM solutions) and incubated (30 min, RT). The samples were centrifuged (10 min, 15000g, RT) and the supernatant was removed. The fibril pellets were washed twice with PBS-T (phosphate buffered saline with 0.1% v/v Tween20). The pellets were resuspended in 35 µL PBS and the fluorescence signal (bound to the fibrils) was measured in a 384-well black microtitre plate (Corning) with a CLARIOstar® Microplate Reader (BMG Labtech, Aylesbury, UK) with an excitation wavelength of 622 nm and collecting the emission spectrum between 650-710 nm. The spectra were corrected for the labelling efficiency of each CTPR-variant and normalised to the 2XFibSyn-CTPR4 readings. Data from two independent experiments were averaged.

### Microfluidic Diffusional Sizing (MDS) experiments

Fabrication and operation of the microfluidic chips for MDS experiments have been shown previously (Arosio et al. 2016; McDonald and Whitesides 2002; Qin et al. 2010). In brief, microfluidic devices were obtained by standard soft-lithography in polydimethylsiloxane (PDMS, Momentive RTV615, Techsil, Bidford on Avon, UK). Carbon nanoparticles (ca. 13 nm, Plasmachem, Berlin, Germany) were mixed into the PDMS before curing to avoid channel cross-talk, as previously reported (Herling et al. 2016). After curation, the PDMS slips were bonded onto microscopy slides (Epredia Cut Microscope Slide, ThermoFisher) by application of an oxygen plasma. Sample and co-flow buffer were loaded onto the chip from reservoirs at the respective inlets by applying a negative pressure at the outlet by means of a glass syringe (Hamilton) connected to a syringe pump (neMESYS, Cetoni GmbH, Korbussen, Germany) with a typical flow rate of 100 µL/min. A custom-built, inverted epifluorescence microscope was used for the detection of the microfluidic profiles. For excitation, light from a brightfield LED light source (Thorlabs, Newton, NJ, USA) was directed through the Cy5-4040C-000 Filter set from Semrock (Laser 2000, Huntingdon, UK) and detected with charge-coupled-device camera (Prime 95B, Photometrics, Tucson, AZ, USA). Images were taken using Micro Manager (Version l.4.23 20170327). Synuclein fibril concentrations are determined by incubation of the fibrils with 4 M GdnHCl and subsequent UV absorption spectroscopy ( χ_275 nm_ = 5600 cm^-1^ M^-1^), and are reported with respect to monomer equivalents. Affinities were determined by determining the hydrodynamic radius of protein complex between the CTPR variant with αS fibrils at different concentrations of both Alexa-647 labelled CTPR variant and unlabelled αS PFFs. With fitting this data with respect to two species, i.e. bound and unbound CTPR variant, it becomes possible to determine the size of the complex along with the fraction of bound CTPR variant. The fraction of bound CTPR variant can be determined with respect to Equation 5 (Scheidt et al. 2021; Scheidt et al. 2019).

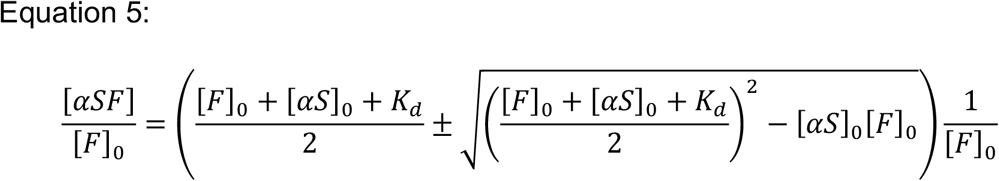

### Fluorescence Microscopy

µ-slide 8 well high glass bottom plates (Ibidi) were treated by cleaning with 1M KOH (30-45 min) and rinsed with PBS (2X). To the well, a poly-L-lysine solution (0.1%) was added and incubated (45 min) followed by washing with PBS (2X). The fibrils bound to Alexa-647 labelled CTPR proteins (10-15 µM) were prepared as described above and these were incubated in a well (30 min), followed by removal of the solution and washing with PBS (1X). Imaging was done on an inverted Olympus IX-73 microscope (Olympus Corporation, Japan), using a TIRF oil-immersion objective (Apochromat 100×, NA 1.49 Oil, Olympus) with an exposer time of 0.01 sec, 100 frames. After being filtered through a dichroic filter cube (Cairn OptoSpin) and a wavelength-specific emission filter (Semrock), images were captured with an electron-multiplying charge-coupled device (EMCCD) camera (Andor iXon3 897) and processed using ImageJ.

## Supplementary Information

The file “Bismut Schneider et al SI 2025.pdf”, contains a table of the amino acid sequences for all designed protein scaffolds, characterization of control protein scaffolds and additional experiments to support the main manuscript.

## Acknowledgements

The authors would like to thank the staff at Cambridge Advanced Imaging Centre (CAIC), University of Cambridge, UK for their assistance with transmission electron microscopy imaging. M.M.S. was funded from the Frances and Augustus Newman Foundation. This work was supported by an MRC Career Development Award (MR/W01632X/1, J.R.K.) and the Royal Society (RGS\R1\231207, J.R.K.). J.R.K. and L.S.I acknowledge funding from the Rosetrees Trust (CF2\100013). G.S.K.S acknowledges funding from the Wellcome Trust (065807/Z/01/Z, 203249/Z/16/Z), the UK Medical Research Council (MR/K02292X/1), Alzheimer’s Research UK (ARUK-PG013-14), the Michael J Fox Foundation (16238, 022159), and Infinitus (China) Company Ltd. T.P.J.K. acknowledges the European Research Council under the European Union’s Horizon 2020 research and innovation program through the ERC grant DiProPhys (agreement ID 101001615).

## Author Contributions

L.S.I. and J.R.K designed the conceptual framework of the study. S.B., M.M.S., M.M., Y.F., E.J.W., M.D.G. and J.R.K. performed the experiments. S.B., M.M.S., M.M., Y.F., E.J.W., M.D.G. T.P.J.K., G.S.K.S, L.S.I and J.R.K. contributed to data acquisition and interpretation. J.R.K wrote the manuscript with contributions from all authors.

