## Supplementary Information for "Using a stable protein scaffold to display peptides that bind to alpha-synuclein fibrils"

**Table S1: Protein sequences and mass spectrometry analysis**

Amino acid sequences for the different CTPR variants show a solvating helix (blue) and an N-terminal His-tag at the N-terminus (italics). Fibril-binding motifs are shown in bold/italics and they are flanked by DPNN linker regions (underlined).

| Protein | Amino acid sequence | Expected Mass (Da) | Observed Mass (Da) |
|---|---|---|---|
| CPTR3 | MRGS*HHHHHH*GLVPRGSAEAWYNLGNAYYKQGDYQKAIEYYQKALELDPRSAEAWYNLGNAYYKQGDYQKAIEYYQKALELDPRSAEAWYNLGNAYYKQGDYQKAIEYYQKALELDPRSAEAKQNLGNAKQKQG | 15633.2 | 15633.2 ± 1.0 |
| Cys-CTPR3 | MRGS*HHHHHH*GLVPRGCAEAWYNLGNAYYKQGDYQKAIEYYQKALELDPRSAEAWYNLGNAYYKQGDYQKAIEYYQKALELDPRSAEAWYNLGNAYYKQGDYQKAIEYYQKALELDPRSAEAKQNLGNAKQKQG | 15649.2 | 15648.6 ± 0.2 |
| FibSyn-CTPR3 | MRGS*HHHHHH*GLVPRGSAEAWYNLGNAYYKQGDYQKAIEYYQKALELDPNN***KLVFFAE***DPNNAEAWYNLGNAYYKQGDYQKAIEYYQKALELDPRSAEAWYNLGNAYYKQGDYQKAIEYYQKALELDPRSAEAKQNLGNAKQKQG | 16893.6 | 16892.6 ± 0.7 |
| Cys-FibSyn-CTPR3 | MRGS*HHHHHH*GLVPRGCAEAWYNLGNAYYKQGDYQKAIEYYQKALELDPNN***KLVFFAE***DPNNAEAWYNLGNAYYKQGDYQKAIEYYQKALELDPRSAEAWYNLGNAYYKQGDYQKAIEYYQKALELDPRSAEAKQNLGNAKQKQG | 16909.6 | 16915.3 ± 6.0 |
| FibSyn2-CTPR3 | MRGS*HHHHHH*GLVPRGSAEAWYNLGNAYYKQGDYQKAIEYYQKALELDPNN***KLVFWAK***DPNNAEAWYNLGNAYYKQGDYQKAIEYYQKALELDPRSAEAWYNLGNAYYKQGDYQKAIEYYQKALELDPRSAEAKQNLGNAKQKQG | 16931.6 | 16931.7 ± 0.1 |
| Cys-FibSyn2-CTPR3 | MRGS*HHHHHH*GLVPRGCAEAWYNLGNAYYKQGDYQKAIEYYQKALELDPNN***KLVFWAK***DPNNAEAWYNLGNAYYKQGDYQKAIEYYQKALELDPRSAEAWYNLGNAYYKQGDYQKAIEYYQKALELDPRSAEAKQNLGNAKQKQG | 16947.7 | 16947.7 ± 0.2 |
| CTPR4 | MRGS*HHHHHH*GLVPRGSAEAWYNLGNAYYKQGDYQKAIEYYQKALELDPNNAEAWYNLGNAYYKQGDYQKAIEYYQKALELDPNNAEAWYNLGNAYYKQGDYQKAIEYYQKALELDPNNAEAWYNLGNAYYKQGDYQKAIEYYQKALELDPNN | 18048.6 | 18054.8 ± 6.7 |
| Cys-CTPR4 | MRGS*HHHHHH*GLVPRGCAEAWYNLGNAYYKQGDYQKAIEYYQKALELDPNNAEAWYNLGNAYYKQGDYQKAIEYYQKALELDPNNAEAWYNLGNAYYKQGDYQKAIEYYQKALELDPNNA | 18064.7 | 18066.5 ± 2.8 |

| | | | |
|---|---|---|---|
| | EAWYNLGNAYYKQGDYQKAIEYYQKALELD PNN | | |
| 2XFibSyn-CTPR4 | MRGSHHHHHHGLVPRGSAEAWYNLGNAYY KQGDYQKAIEYYQKALELDPNN***KLVFFAE***DP NNAEAWYNLGNAYYKQGDYQKAIEYYQKAL ELDPNNAEAWYNLGNAYYKQGDYQKAIEYY QKALELDPNN***KLVFFAE***DPNNAEAWYNLGN AYYKQGDYQKAIEYYQKALELDPNN | 20599.5 | 20599.0 ± 2.0 |
| Cys-2XFibSyn-CTPR4 | MRGSHHHHHHGLVPRGCAEAWYNLGNAYY KQGDYQKAIEYYQKALELDPNN***KLVFFAE***DP NNAEAWYNLGNAYYKQGDYQKAIEYYQKAL ELDPNNAEAWYNLGNAYYKQGDYQKAIEYY QKALELDPNN***KLVFFAE***DPNNAEAWYNLGN AYYKQGDYQKAIEYYQKALELDPNN | 20615.6 | 20615.6 ± 0.2 |

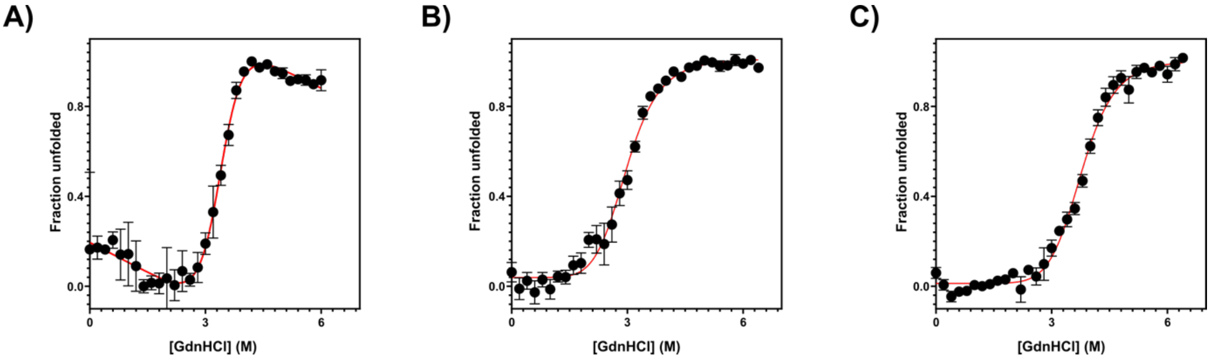


**Fig. S1. GdnHCl-induced denaturation of CTPR proteins.** Purification of CTPR variants yielded samples with concentrations between 150-400 μM. A) FibSyn-CTPR3, B) FibSyn2-CTPR3 and C) 2XFibSyn-CTPR4CTPR4. 1 μm of protein in 50 mM sodium phosphate, 150 mM NaCl, pH 7.5 and increasing GdnHCl concentrations (from 0 M to 6.4 M) at 25°C. Excitation was at 295 nm and emission intensity was measured at 360 nm. A minimum of three independent experiments were recorded for each protein. The data were fitted with a two-state model to give the midpoint of unfolding and *m* value (a constant proportional to the change in solvent-accessible surface area upon unfolding). Error bars show the standard error of the mean.

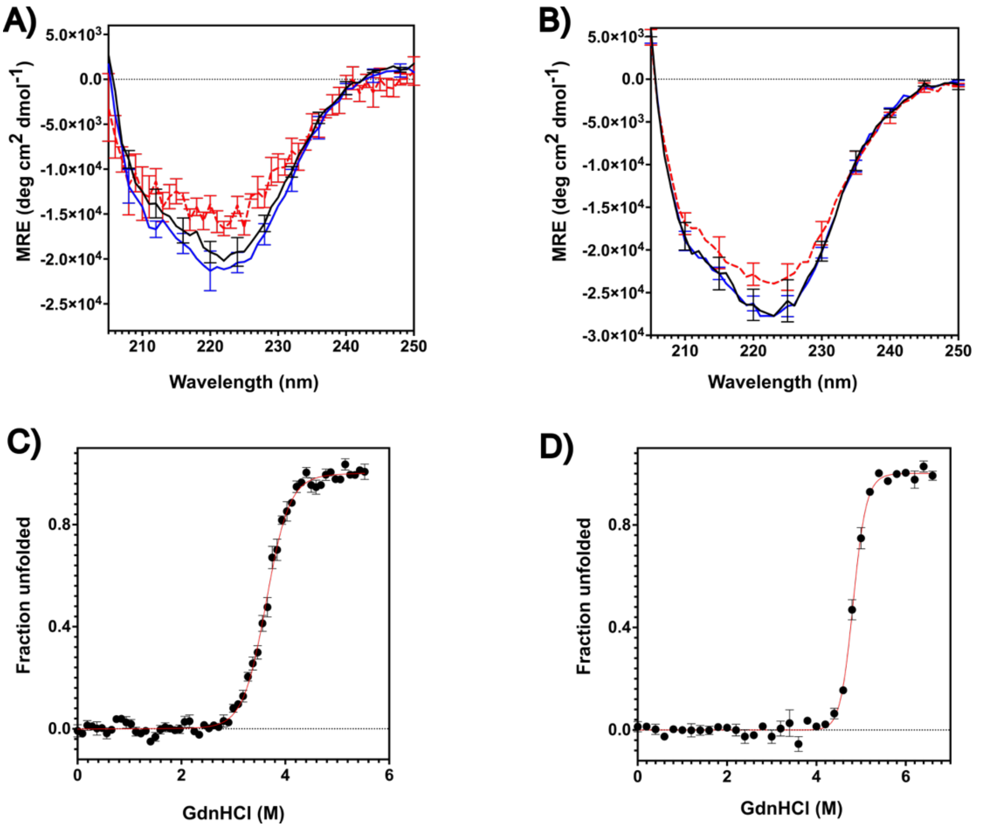


**Fig. S2. CD analysis and GdnHCl-induced denaturation of CTPR3 and CTPR4 (negative controls).** To determine whether fibril-binding is due to the presence of the peptide motifs, CTPR3 and CTPR4 containing the native inter-repeat loops were used. CD analysis of A) CTPR3 and B) CTPR4 was performed with 5 µm of protein in 50 mM sodium phosphate, 150 mM NaCl, 0.5 mM TCEP, pH 7.5, spectra recorded from 205 nm to 250 nm in 0.5 nm intervals. Samples were measured at 20°C (black), 90°C (red) and return to 20°C (blue). GdnHCl denaturation was performed for C) CTPR3 and D) CTPR4 with 1 µm of protein in 50 mM sodium phosphate, 150 mM NaCl, pH 7.5 and increasing GdnHCl concentration (from 0 M to 6.4 M) at 25°C. Excitation was at 295 nm and emission intensity was measured at 360 nm. A minimum of three independent experiments were recorded for each protein. The data were fitted to a two-state model to give the midpoint of unfolding and *m* value (a constant proportional to the change in solvent-accessible surface area upon unfolding). Error bars show the standard error of the mean

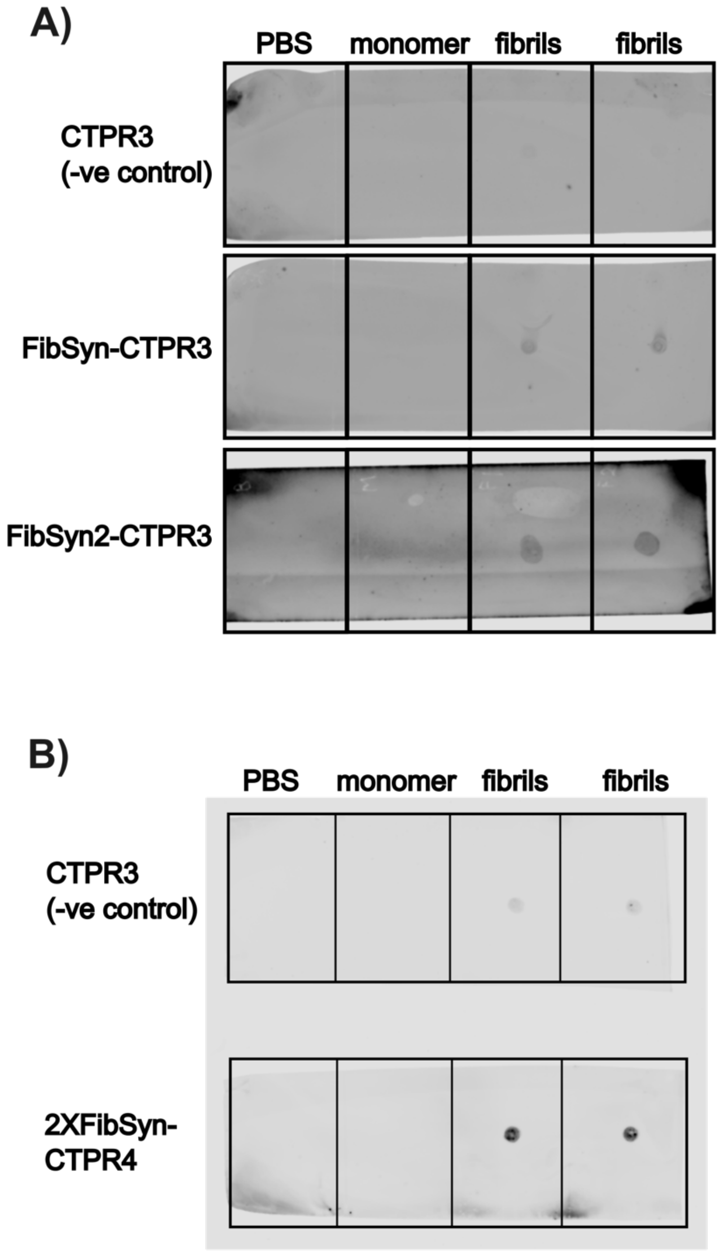


**Fig. S3. Dot blot assay to test CTPR variant binding to monomeric and fibrillar $\alpha$-synuclein.** Monomeric and fibrillar $\alpha$-synuclein samples were applied to nitrocellulose membranes followed by incubation with A) Alexa Fluor$^{\mathrm{TM}}$ 594-labelled CTPR variants and B) Alexa Fluor$^{\mathrm{TM}}$ 647-labelled CTPR variants. Imaging using a LiCor Odessey system shows no interactions between the fluorophore-labelled CTPRs and monomeric $\alpha$-synuclein and increased interactions between fibrillar $\alpha$-synuclein and CTPR variants containing the fibril-binding peptide motifs. Some non-specific binding was observed with the Alexa Fluor$^{\mathrm{TM}}$ 647-labelled CTPR3 negative control (also observed in the fibril-pulldown assay), this non-specific binding was not observed with the Alexa Fluor$^{\mathrm{TM}}$ 594-labelled CTPR3 negative control.

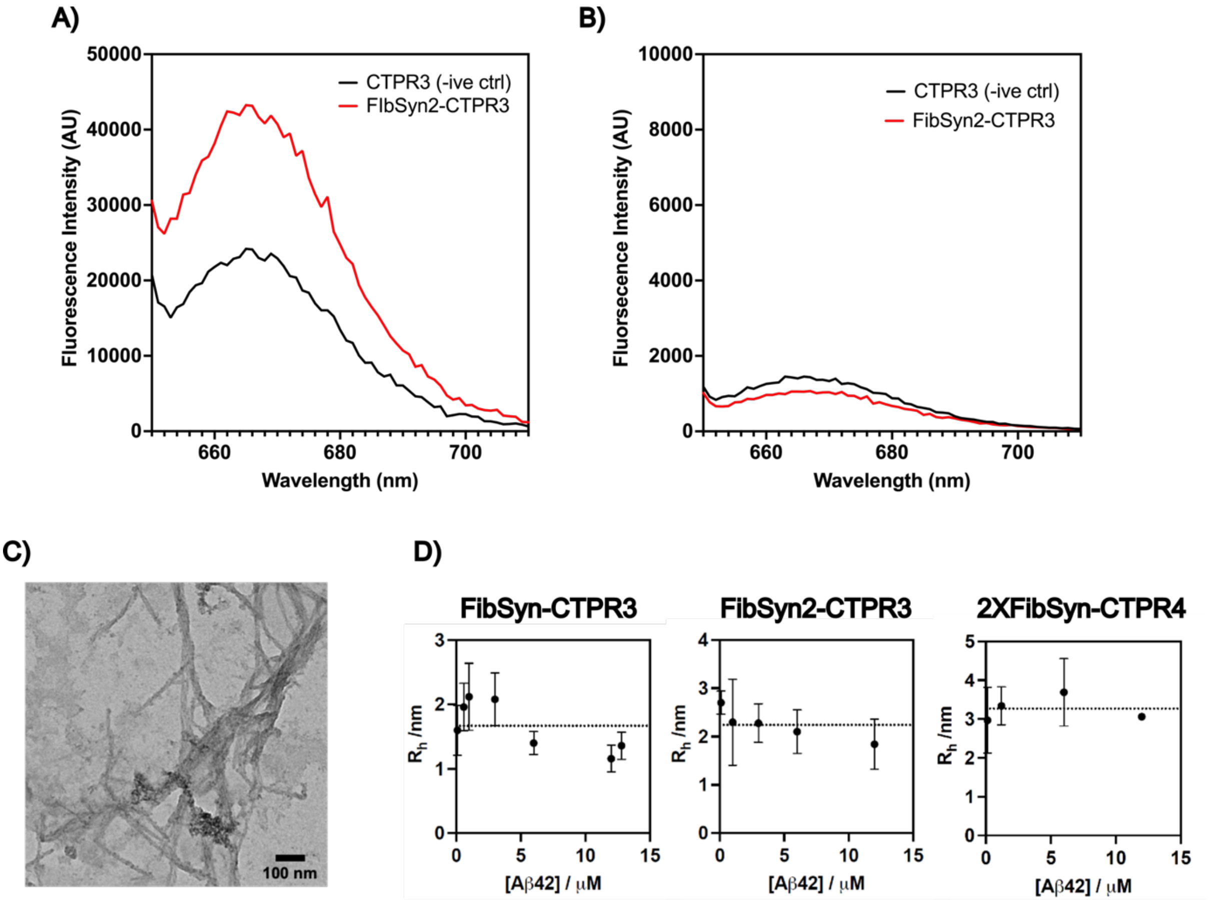


**Fig S4. Comparing CTPR variants interactions with $\alpha$-synuclein fibrils versus $A\beta_{1-42}$ fibrils.** A) Fluorescence emission spectrum for CTPR3 (negative control) and FibSyn-CTPR3 bound $\alpha$-synuclein fibrils showing specific interactions. B) Fluorescence emission spectrum for CTPR3 (negative control) and FibSyn-CTPR3 bound $A\beta_{1-42}$ fibrils, showing no specific interactions with FibSyn-CTPR3. C) Representative TEM image of $A\beta_{1-42}$ fibrils used for fibril pull-down and MDS assays. D) MDS assays of Alexa Fluor$^{TM}$ 647-labelled CTPR variants with increasing concentrations of $A\beta_{1-42}$ fibrils. No change in hydrodynamic radius is seen for any of the FibSyn-CTPRs.
